# Ras-Responsive Element Binding Protein 1 regulates survival of Group 3 medulloblastoma

**DOI:** 10.1101/2025.07.16.665230

**Authors:** Meher Beigi Masihi, Kendall R. Chambers, Richard A. Friedman, Sergey Pampou, Brian L. Gudenas, Jacob Torrejon, Catherine Lee, Grace Furnari, Lianne Q. Chau, Sajina GC, Yingxi Lin, Owen S. Chapman, Patryk Skowron, Livia Garzia, Michael D. Taylor, Lukas Chavez, Olivier Ayrault, Paul A. Northcott, Charles Karan, Robert J. Wechsler-Reya

## Abstract

Medulloblastoma (MB) is the most common malignant pediatric brain tumor, and Group 3 (G3, MYC-driven) MB has the worst prognosis. Despite advances in molecular classification, oncogenic drivers of G3 MB remain poorly defined. To identify such drivers, we profiled transcription factor expression across MB subgroups. Our analysis revealed Ras-responsive element binding protein 1 (RREB1) as one of the most highly expressed transcription factors in G3 MB. RREB1 knockdown impaired cell proliferation in vitro and prolonged survival in orthotopic xenograft models, suggesting it plays a key role in regulating tumor growth. Mechanistically, RREB1 acts by enhancing transcription of TGF-β pathway genes. Upstream, RREB1 expression is controlled by c-MET signaling, and the MET inhibitor SU11274 decreased RREB1 levels and MB cell viability. Local delivery of SU11274 in tumor-bearing mice suppressed RREB1 expression and extended survival. These results establish the c-MET/RREB1 axis as a critical oncogenic regulator and a promising therapeutic target in in high-risk MB.

## Introduction

Medulloblastoma (MB) is the most common malignant childhood brain tumor (1,2). Although surgery, radiation therapy and chemotherapy have significantly improved survival, ∼1/3 of patients still die from the disease, and survivors suffer severe long-term side effects from therapy (3,4). Therefore, more effective and less toxic therapies are desperately needed.

MB is a heterogenous disease, and recent molecular analysis suggests that it can be divided into four major subgroups – WNT, Sonic Hedgehog (SHH), Group 3 (G3-MB), and Group 4 (G4-MB). Each subgroup is characterized by distinct genetic and epigenetic profiles and clinical outcomes (5,6). Among these subgroups, G3-MB accounts for ∼25% of MB patients and has the poorest prognosis (7). Nearly half of G3-MB patients present with metastases at the time of diagnosis, representing the highest incidence among all subgroups. Unlike WNT and SHH MBs, there are no known germline mutations that predispose to G3-MB. However, somatic copy number alterations are frequently observed in these tumors. Among them, amplification of the MYC oncogene stands out prominently, affecting approximately 17% of G3-MB patients (7–9). MYCN, a member of the MYC family transcription factors, is frequently amplified in SHH and G4-MB, where its overexpression correlates with unfavorable clinical outcomes (6,10). The transcription factor OTX2 (orthodenticle homeobox 2) is another key driver in G3 and G4-MB, with its overexpression generally mutually exclusive of MYC amplification (11,12).

Given the importance of transcription factors in regulating cell fate, proliferation, survival and differentiation, we sought to identify novel transcription factors that might function as drivers of G3-MB. By analyzing MB transcriptomic data we uncovered a transcription factor, Ras-responsive element binding protein 1 (RREB1), that is highly expressed in G3-MB. Here we evaluate the role of RREB1 in proliferation and survival, explore the signals that regulate its expression and the targets that mediate its effects, and examine the implications of these pathways for therapy of this disease.

## Results

### Integrated gene expression analysis reveals overexpression of RREB1 in G3 and G4 medulloblastoma cells

To identify transcription factors differentially expressed in Group 3 medulloblastoma (G3-MB), we analyzed a publicly available transcriptomic dataset of 763 human MB tumors classified into four molecular subgroups (WNT, *n* = 70; SHH, *n* = 223; G3, *n* = 144; G4, *n* = 326) (10,13).

Focusing on genes encoding DNA-binding transcription factors, we identified RREB1 as one of the most significantly upregulated candidates in G3- and G4-MB relative to WNT and SHH subgroups (Fig. 1a-c; Supplementary Table 1). To validate these findings, we examined RREB1 expression in an independent transcriptomic dataset comprising 341 MB samples. Consistent with the previous finding, RREB1 mRNA levels were elevated in G3 and G4 tumors compared to WNT and SHH subtypes (Fig. 1d). Notably, RREB1 expression was minimal in normal prenatal and adult brain tissues (Supplementary Fig. 1a), suggesting tumor-specific activation.

**Figure 1.**
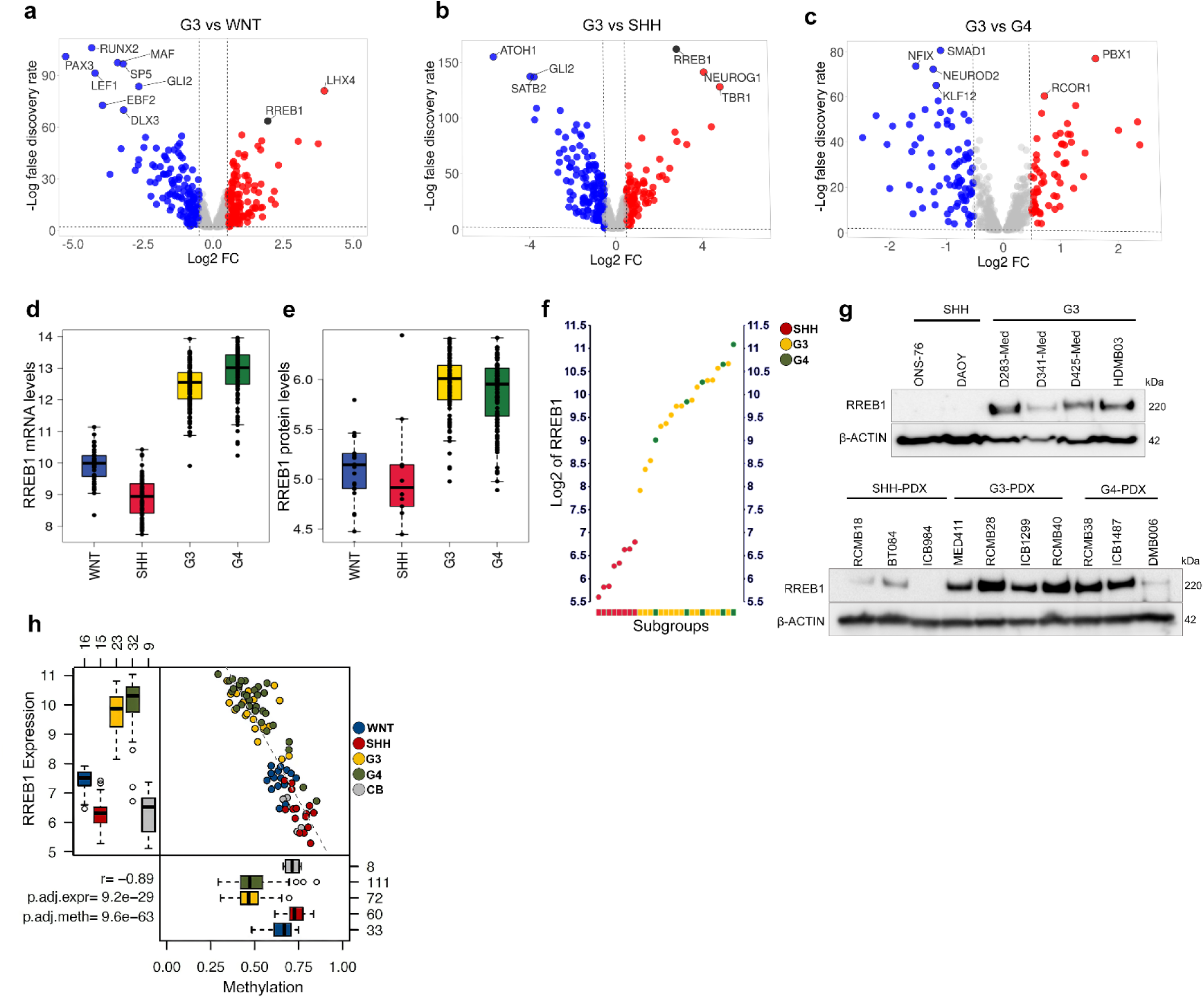
RREB1 is overexpressed in G3 and G4-MB. **a-c.** Differential expression of DNA-binding transcription factors across medulloblastoma (MB) subgroups (n = 763; WNT = 70, SHH = 223, Group 3 = 144, Group 4 = 326; data from Cavalli et al. obtained via the R2 Genomic Analysis and Visualization Platform (R2), https://hgserver1.amc.nl/). Differentially expressed genes were selected based on an absolute log2 fold change ≥ 1 and a false discovery rate < 0.01. **d.** RREB1 mRNA expression is significantly elevated in Group 3 and Group 4 medulloblastoma compared to WNT and SHH subgroups (*n* = 341). **e.** RREB1 protein expression, assessed by proteomic analysis, shows a similar pattern of upregulation in Group 3 and Group 4 tumors (*n* = 343). **f.** RREB1 mRNA expression levels in 27 patient-derived xenografts (PDXs) representing SHH, G3, and G4 MB obtained from R2. **g.** Western blot analysis showing elevated protein levels in G3 and G4-MB PDXs compared to SHH. Protein lysates from G3, G4, and SHH-MB samples were subjected to SDS-PAGE and probed with an antibody against RREB1. β-ACTIN was used as a loading control. **h.** An inverse correlation between RREB1 promoter methylation (assessed at 21 CpG sites using the Illumina 450k array in 42 MB samples across all subgroups) and mRNA expression (measured with the Affymetrix U133 Plus 2.0 array in 95 MB samples) was observed. Group 3/4 tumors exhibit hypomethylation, indicative of an active promoter and high RREB1 expression, whereas WNT/SHH tumors display hypermethylation and low expression (Pearson’s r = −0.89; expression adj. p = 9.2e-29; methylation adj. p = 9.6e-63). CB indicates healthy cerebellum.

Proteomic analyses in a cohort of 343 MB samples demonstrated elevated RREB1 protein levels in G3- and G4-MB (Fig. 1e), while evaluation of MB cell lines and patient-derived xenografts (PDXs) showed RREB1 to be highly overexpressed in G3/G4 tumors at both the RNA (Fig. 1f, Supplementary Fig. 1b), and protein level (Fig. 1g). Together, these data indicate that RREB1 is highly expressed in G3 and G4-MB.

To investigate the molecular basis for increased expression of RREB1 in G3 and G4, we analyzed publicly available array-based DNA methylation data (14,15). Our studies revealed a notable anticorrelation between RREB1 expression and CpG methylation proximal to the RREB1 promoter (n=21 CpGs) across all MB subgroups. G3 and G4 tumors exhibited the lowest methylation levels and the highest expression levels, suggesting an active (unmethylated) RREB1 promoter in these tumors (Fig. 1h). In contrast, SHH and WNT tumors displayed much stronger methylation and lower expression, consistent with the repression of gene expression at the RREB1 locus. These results suggest that overexpression of RREB1 in G3/G4 tumors is associated with the presence of active promoters due to minimal methylation.

### RREB1 is required for tumor growth in G3-MB

The heightened expression of RREB1 in G3-MB prompted us to examine its role in tumor growth. To this end, we used lentiviral short hairpin RNAs (shRNAs) co-expressing green fluorescent protein (GFP) to knock down RREB1 expression in the human G3-MB cell lines D283-Med and D341-Med, which exhibit elevated levels of RREB1 expression (Supplementary Fig. 1b). Knockdown efficiency was confirmed by sorting infected cells for GFP expression and examining protein levels by Western blotting 48-hours post-transduction (Supplementary Fig. 1c).

To examine the impact of RREB1 knockdown on cell proliferation, we performed BrdU incorporation assays. These studies revealed a significant reduction in BrdU incorporation in G3-MB cell lines 96-hours after RREB1 knockdown (Fig. 2a). To characterize the mechanism underlying the diminished viability, we assessed apoptosis and cell death by flow cytometry using annexin V and 7-AAD staining. RREB1 knockdown cells exhibited a marked increase in annexin V and 7-AAD double-positive cells compared to controls (Fig. 2b), indicating progression to late apoptosis and cell death. CellTiter-Glo viability assays conducted over a span of 10 days also demonstrated a progressive reduction in viable cell number in G3 cell lines following RREB1 knockdown (Fig. 2c, d). Collectively, these findings demonstrate that loss of RREB1 impairs cell proliferation and induces apoptotic cell death in G3-MB cells.

**Figure 2.**
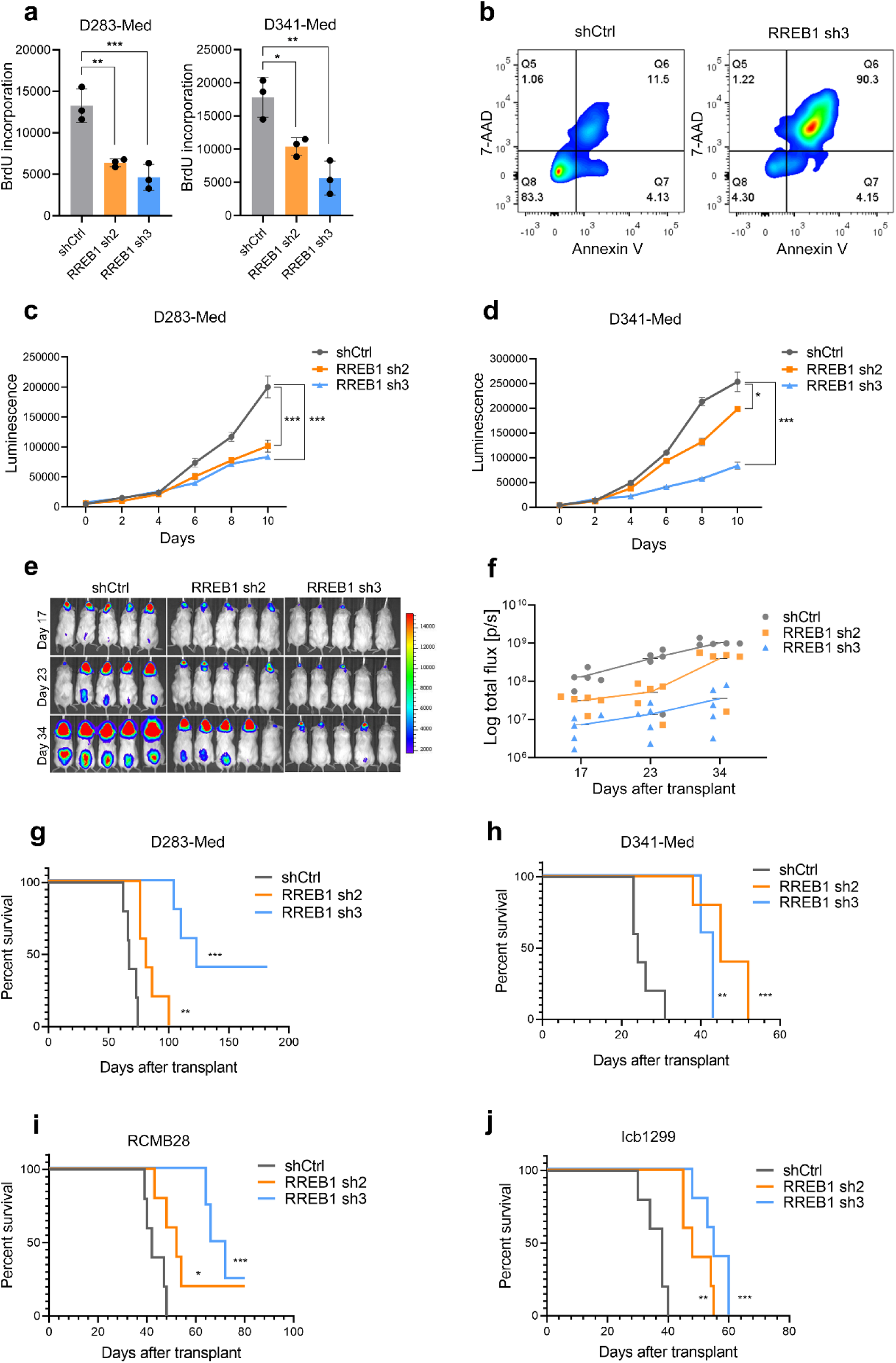
RREB1 is required for the growth of G3-MB cells. **a.** Quantification of cell proliferation using a colorimetric BrdU assay at 96-hours post-knockdown. Absorbance was measured at 450 nm. Statistical analysis was performed using one-way ANOVA; error bars represent mean ± SD from three biological replicates (* p < 0.05, ** p < 0.01, *** p < 0.001). **b.** Flow cytometric analysis of apoptosis after RREB1 knockdown. Cells were sorted and cultured for 5 days prior to staining with Annexin V (apoptotic cells) and 7-AAD (dead cells). Cells were incubated in Annexin V binding buffer at a 1:100 dilution and 7-AAD at a 1:100 dilution for 15 minutes before analysis. **c, d.** Longitudinal assessment of cell viability using the CellTiter-Glo assay. Infected cells were sorted at 48 hours post-transduction, plated, and luminescence were measured every other day. Each data point represents the mean ± SD from three independent experiments, with significance determined by unpaired t-test (* p < 0.05, *** p < 0.001). **e.** In vivo bioluminescent imaging (IVIS) of mice implanted with RREB1 knockdown or control cells. Mice received an intraperitoneal injection of 75 mg/kg luciferin and were imaged on days 17, 23, and 34. **f.** Bioluminescence is expressed as Log10 total flux (photons/second) (n = 5 per group). **g, h.** In vivo tumor growth and survival were evaluated in naïve NSG mice implanted with RREB1 knockdown cells. The median survival times for the D283-Med group were 67 days for shCtrl, 81 days for RREB1 sh2, and 123 days for RREB1 sh3; for the D341-Med group, median survival was 24, 45, and 43 days, respectively. i**, j.** The median survival for RCMB28 was 42 days for shCtrl, 52 days for RREB1 sh2, and 69 days for RREB1 sh3; and for the ICB1299 group, the median survival was 38, 48, and 55 days respectively. Kaplan-Meier survival curves were generated and statistically compared using the Log-rank (Mantel-Cox) test (n = 5 per group, * p < 0.05, ** p < 0.01, *** p < 0.001).

To assess the impact of these effects on tumor growth in vivo, RREB1 was knocked down in D283-Med and D341-Med cell lines expressing firefly luciferase (F-luc). Cells were transplanted into the cerebellum of immunodeficient (NOD-SCID-IL2Rgamma knockout, or NSG) mice, and animals were monitored for tumor growth via bioluminescence imaging. As shown in Fig. 2e-h mice transplanted with D283-Med cells or D341-Med cells in which RREB1 had been knocked down exhibited prolonged survival compared to those receiving cells infected with scrambled shRNA (shCtrl). Similar results were seen with the G3 MB PDXs RCMB28 and Icb1299, which express high levels of RREB1 (Fig. 2i, j). These results strongly suggest that RREB1 knockdown not only inhibits cell proliferation in vitro but also confers a survival advantage in vivo, underscoring its potential as a therapeutic target in G3-MB.

### RREB1 regulates TGF-β pathway target genes

To understand the mechanisms underlying the effects of RREB1 on proliferation, we generated D283-Med cells expressing a doxycycline (DOX)-inducible RREB1 shRNA. Treatment of these cells with DOX induced robust knockdown of RREB1 at 24 and 48-hours (Supplementary Fig. 2a). We then conducted RNA sequencing (RNA-seq) on RREB1-shRNA expressing cells 24-hours post-DOX induction, utilizing cells expressing a DOX-inducible scrambled shRNA as controls. Differential expression analysis identified 1015 genes exhibiting a log2 fold change greater than 0.6 or less than −0.6 and a p-value less than 0.001 in RREB1 knockdown cells compared to controls (Fig. 3a). KEGG Pathway analysis showed enrichment of pathways associated with TGF-β signaling, neuroactive ligand-receptor interaction, cytokine-cytokine interaction, and calcium signaling pathways (Fig. 3b). Notably, the majority of the TGF-β pathway target genes were found to be downregulated following RREB1 knockdown, as confirmed by quantitative RT-PCR (Fig. 3c, d). To determine whether RREB1 modulates TGF-β pathway targets by altering activity of SMAD proteins, we infected D283-Med or D341-Med cells with a lentiviral construct containing SMAD2/3/4 binding motifs upstream of a F-luc reporter. RREB1 knockdown resulted in a significant suppression of Activin B-induced, but not TGF-β-induced, SMAD activity when compared to cells infected with shCtrl viruses (Fig. 3e, f). Given the critical role of the TGF-β pathway in cell cycle and differentiation in MB and other cancers, (16–18), we hypothesized that RREB1 might mediate its effects in part through this pathway.

**Figure 3.**
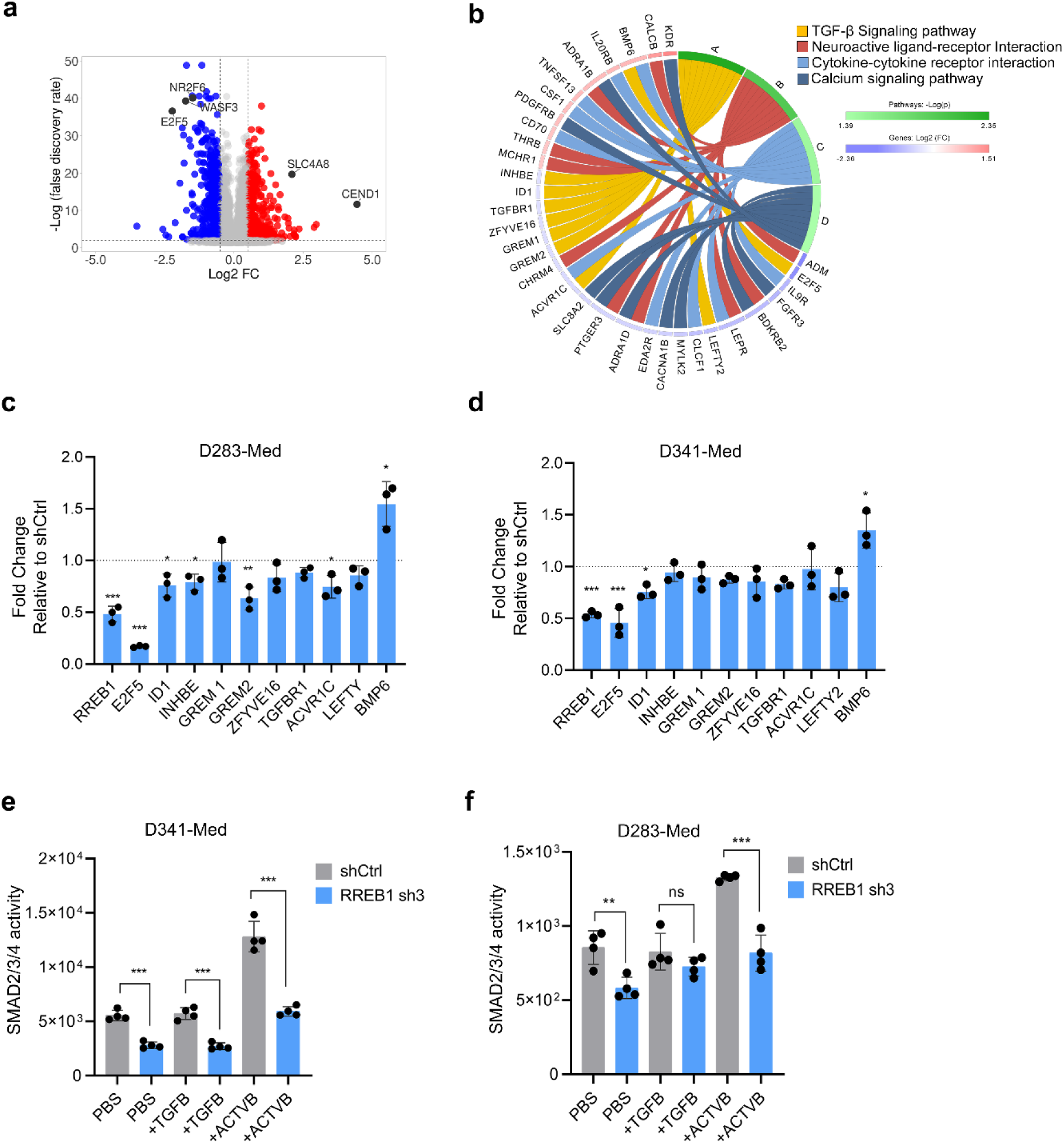
RREB1 regulates TGF-β target genes in G3-MB cells. **a.** The volcano plot illustrates the significantly altered genes in D283-Med cells transduced with shRNA3 targeting RREB1 compared to shCtrl. Differential expression was defined by a p-value < 0.01, fdr < 0.05, and an absolute log2 fold change > 0.6, based on three independent biological replicates. In total, 428 genes were significantly downregulated, whereas 402 genes were significantly upregulated following RREB1 knockdown. **b.** Circus plot illustrating the enriched pathways identified among the differentially expressed genes. **c, d.** qPCR validation of TGF-β pathway-associated genes upon RREB1 knockdown in D283-Med and D341-Med cells, respectively. The most significant reduction was associated with E2F5 expression (p < 0.001) in both cell lines. Fold changes were calculated using the ΔΔCt method with shCtrl as the normalization control. Data represent the mean ± SD from three independent biological replicates, with statistical significance assessed using a t-test (* p < 0.05, ** p < 0.01, *** p < 0.001). **e, f.** SMAD2/3/4 reported assay. D283-Med and D341-Med cells transduced with RREB1 sh3 or shCtrl were treated with 200 ng/ml TGF-β, Activin B, or PBS (vehicle control). SMAD2/3/4 reporter activity was quantified by luminescence 24-hours post-treatment. Data represent mean ± SD from four independent experiments (unpaired t-test: ***p < 0.001, **p < 0.01, ns = non-significant vs. shCtrl).

The above studies identified changes in gene expression following knockdown of RREB1, but these could include both direct and indirect targets of this transcription factor. To determine which genes are directly bound by RREB1, we overexpressed an HA-tagged human RREB1 (HA-RREB1) coding sequence in D283-Med cells and performed CUT&RUN analysis. This approach revealed 5,721 RREB1 peaks across the genome within 5 kilobases of transcription start sites (TSS) (Fig. 4a). Integration of the RREB1 binding sites from CUT&RUN analysis with the downregulated genes in the TGF-β pathway identified 4 genes – *E2F5*, *ID1*, *INHBE*, and *GREM2* – with altered expression following RREB1 knockdown (Fig. 4b-d). Among these genes, *E2F5* stood out due to its known role in regulating the cell cycle, especially the G1/S transition (19,20).

**Figure 4.**
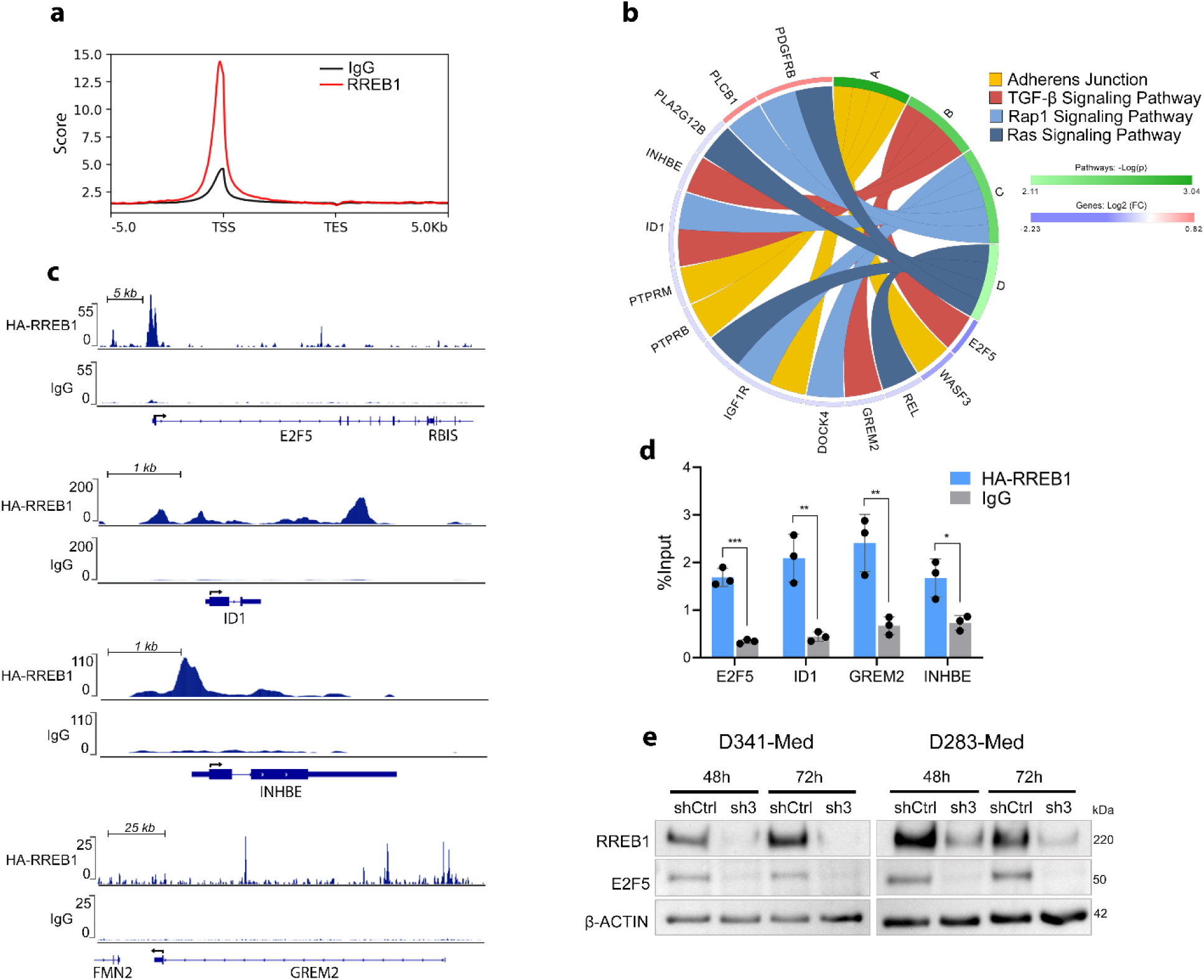
CUT&RUN analysis reveals RREB1 binding to TGF-β target genes. **a.** Genome-wide distribution of RREB1 binding relative to transcription start sites (TSS) and transcription end sites (TES). The plot represents the normalized coverage of RREB1 CUT&RUN signal (red) and IgG control (black) across all annotated genes, with signals averaged over a ±5 kb window around the TSS and TES. **b.** Pathway enrichment analysis of RREB1 direct target genes. Significantly enriched pathways (Adherens junction, TGF-β signaling, Rap1 signaling pathway, and Ras signaling pathway) are ranked by fold enrichment. **c.** CUT&RUN signal tracks at RREB1 target loci. RREB binding at the promoter regions of *E2F5*, *ID1*, *INHBE*, and *GREM2* genes in D283-Med cells. HA-RREB1 CUT&RUN signal (top track, anti-HA antibody) and IgG control (bottom track) across genomic regions of interest. Gene annotations (exons, introns, transcriptional orientation) are displayed below signal tracks. Tracks scaled equally within each panel. Arrows indicate the start of the coding sequence. **d.** CUT&RUN-qPCR validation of RREB1 binding to TGF-β-associated target loci. Chromatin from D283-Med cells was immunoprecipitated with anti-HA (RREB1) or IgG (negative control) antibodies. Enrichment of *E2F5*, *ID1*, *INHBE*, and *GREM2* loci was quantified by qPCR (mean ± SD; n = 3 independent experiments; unpaired t-test, ***p < 0.001, **p < 0.01, *p < 0.05 vs. IgG). **e**. RREB1 knockdown reduces E2F5 protein levels in G3-MB cells. Western blot analysis of E2F5 expression in D283-Med and D341-Med cells transduced with either RREB1 sh3 or shCtrl. Cell lysates were collected at 48- and 72-hours post-transduction, resolved by SDS-PAGE, and immunoblotted with an anti-E2F5 antibody. β-ACTIN was used as a loading control.

Consistent with its identification as a direct RREB1 target, we observed that E2F5 protein levels were reduced upon RREB1 knockdown at both 24 and 48-hours, further confirming that E2F5 is directly regulated by RREB1 (Fig. 4e).

To explore its functional relevance in G3-MB, we employed shRNAs or a pharmacological inhibitor to inhibit E2F5 in MB cell lines. Our studies demonstrated a significant reduction in cell viability in E2F5 knockdown cells compared to control cells expressing shCtrl (Supplementary Fig. 2b). Similarly, the small molecule E2F5 inhibitor HLM006474 (21) significantly impaired MB cell viability (Supplementary Fig. 2c). These findings suggest that RREB1 may act as a transcriptional regulator of TGF-β pathway targets, potentially influencing key cell cycle mediators such as E2F5.

### High-throughput screening identifies c-MET as a regulator of RREB1

While RREB1 has been shown to function as a critical driver of tumor progression, no direct pharmacological inhibitors of this transcription factor currently exist. To identify RREB1-modulating compounds, we employed CRISPR-Cas9 genome editing to tag the N-terminus of the RREB1 gene with a HiBiT luminescent reporter in HDMB03 cells (Fig. 5a). This system enables real-time quantification of RREB1 protein via HiBiT signal intensity. Using this platform, we performed high-throughput screening (HTS) using a 378-compound kinase inhibitor library, treating cells for 8-hours with each compound at 10 µM. The most potent inhibitors of HiBiT luminescence (RREB1 expression) included modulators of BCR-ABL, c-MET, and mTOR signaling nodes (Fig. 5b). Follow-up validation identified the c-MET inhibitor SU11274 as a selective and potent suppressor of RREB1 protein expression (Fig. 5c). Notably, SU11274 caused a dose-dependent reduction in viability of multiple MB cell lines (Fig. 5d-g). Conversely, treatment with hepatocyte growth factor (HGF), a canonical c-MET ligand, increased RREB1 protein levels as well as inducing phosphorylation of ERK but not AKT (Fig. 5h-j). The induction of RREB1 by HGF stimulation and the reduction of RREB1 by c-MET inhibition confirms that RREB1 is a downstream target of the MET pathway. Notably, we further validated that another c-MET inhibitor, foretinib, also reduced RREB1 protein levels in multiple MB cell lines (Supplementary Fig. 3a, b), supporting a role for c-MET signaling in modulating RREB1 stability or expression. Cells also showed sensitivity to foretinib, as determined by IC50 measurements (Supplementary Fig. 3c-e). These findings position c-MET inhibitors as a promising therapeutic strategy for targeting RREB1-driven MB, demonstrating their capacity to suppress RREB1 protein levels at clinically achievable doses.

**Figure 5.**
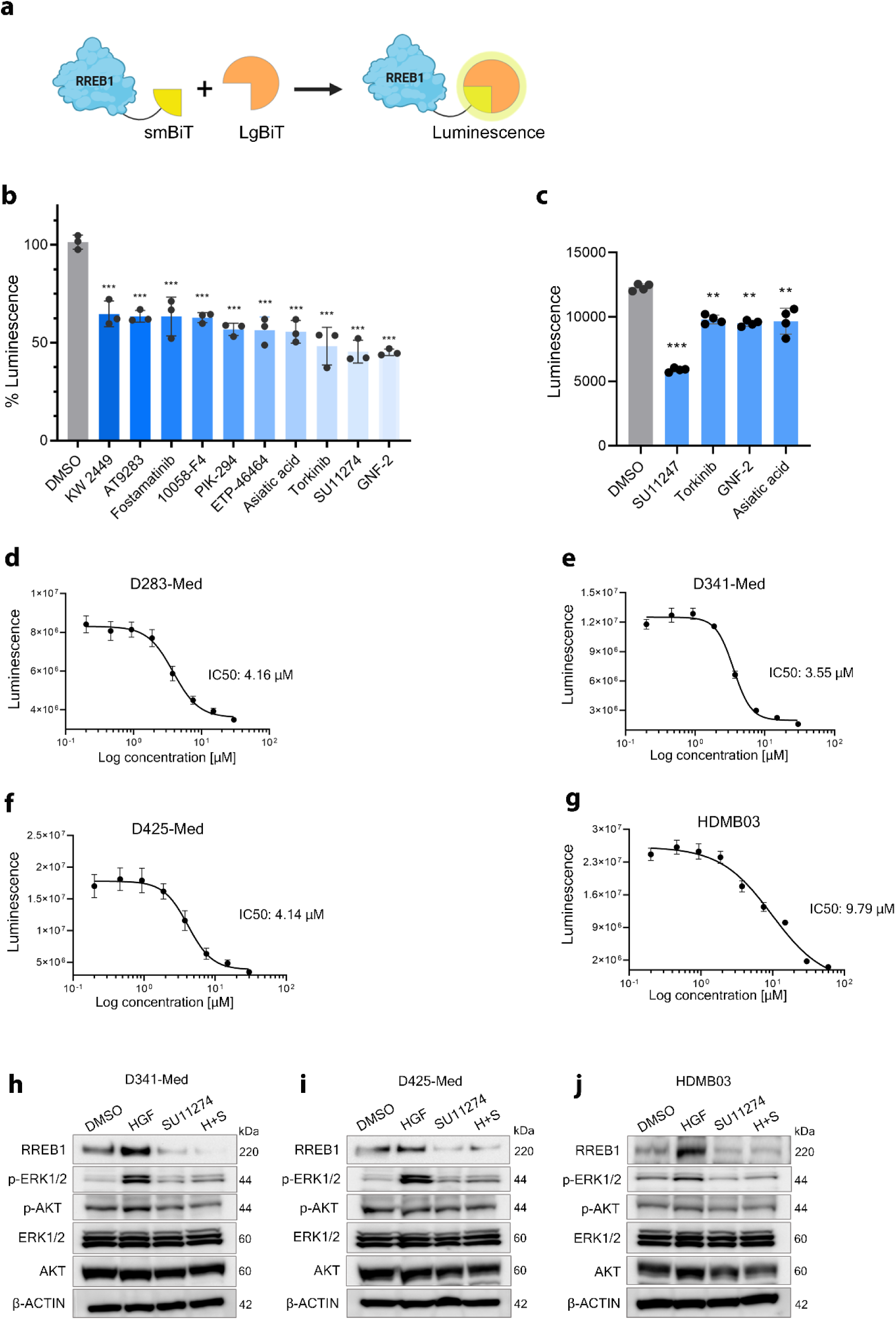
Inhibition of c-MET signaling reduces RREB1 protein levels. **a.** Schematic representation of endogenous HiBiT tagging of RREB1 in HD-MB03 cells. CRISPR– Cas9–mediated genome editing was employed to insert the small BiT (smBiT) tag immediately upstream of the RREB1 coding sequence (N-terminus) via a repair template harboring smBiT flanked by homology arms. Successful integration yields RREB1–smBiT fusion proteins expressed under the native promoter. Upon addition of exogenous large BiT (LgBiT), complementation of smBiT and LgBiT restores NanoLuc luciferase activity, producing a luminescent signal proportional to endogenous RREB1 protein levels. **b.** Top 10 compounds from HTS ranked by relative luminescent reduction (10 µM, 8-hour treatment). GNF-2 exhibited the strongest inhibition (45.23%), followed by SU11274 (45.49%), Torkinib (48.30%), and Asiatic acid (55.67%) versus DMSO control (mean ± SD; n = 3; one-way ANOVA, ***p < 0.001). **c.** Validation of top 5 HTS hits in HiBiT-HDMB03 cells (10 µM, 8 hours; mean ± SD; n = 3; one-way ANOVA, **p < 0.01, ***p < 0.001 vs. DMSO). **d-g.** Dose-response curves of SU11274 in D283-Med (IC50 = 4.16 µM), D341-Med (IC50 = 3.55 µM), D425-Med (IC50 = 4.14 µM), and HDMB03 (IC50 = 9.79 µM) cells. Cells were treated with serially diluted SU11274 (0.1–100 µM) for 72-hours, and luminescence were measured using CellTiter-Glo. Data were log-transformed and fitted to a four-parameter nonlinear regression model (mean ± SD; n = 3 biological replicates). **h-j.** Western blot analysis of RREB1, ERK1/2, phospho-ERK (p-ERK), AKT, and phospho-AKT (p-AKT) in G3-MB cell lines. Cells were serum-starved for 24 hours, followed by treatment with DMSO (control), 200 ng/mL HGF, 4 µM SU11274, or SU11274 + HGF for 48-hours. β-ACTIN served as a loading control. SU11274 attenuated RREB1 protein level and HGF induced RREB1 protein and phosphorylation of ERK but not AKT. H+S: HGF+SU11274.

### Convection-Enhanced Delivery of a c-MET inhibitor prolongs survival of tumor bearing mice

Despite its therapeutic potential, systemic administration of SU11274 did not result in a change in RREB1 levels, suggesting that the drug does not effectively cross the blood-brain barrier (BBB) and accumulate in brain tumors following systemic administration (Supplementary Fig. 4a). To circumvent this challenge, convection-enhanced delivery (CED) — a method enabling continuous, localized infusion of therapeutic agents directly into target tissues — was employed. By bypassing the BBB, CED enhances drug distribution within the tumor microenvironment, offering a strategic advantage for delivering small-molecule inhibitors like SU11274 with heightened precision. In this study, we leveraged CED via osmotic pumps to assess the efficacy of the c-MET inhibitor SU11274 in an orthotopic tumor model, thereby overcoming its pharmacokinetic constraints and optimizing its delivery.

To determine whether intratumoral delivery of SU11274 could reduce RREB1 levels, we employed osmotic pumps with a constant flow rate of 1 µL/hour, allowing for continuous drug release. Pumps were implanted within tumors, and after three days, tumors were harvested for analysis of RREB1 levels, the apoptotic marker cleaved caspase-3 (CC3), and the proliferation marker Ki67 (Fig. 6a-e; Supplementary Fig. 4b). SU11274 treatment resulted in a marked reduction in RREB1and Ki67 expression accompanied by a robust increase in CC3, consistent with induction of apoptosis in tumor cells.

**Figure 6.**
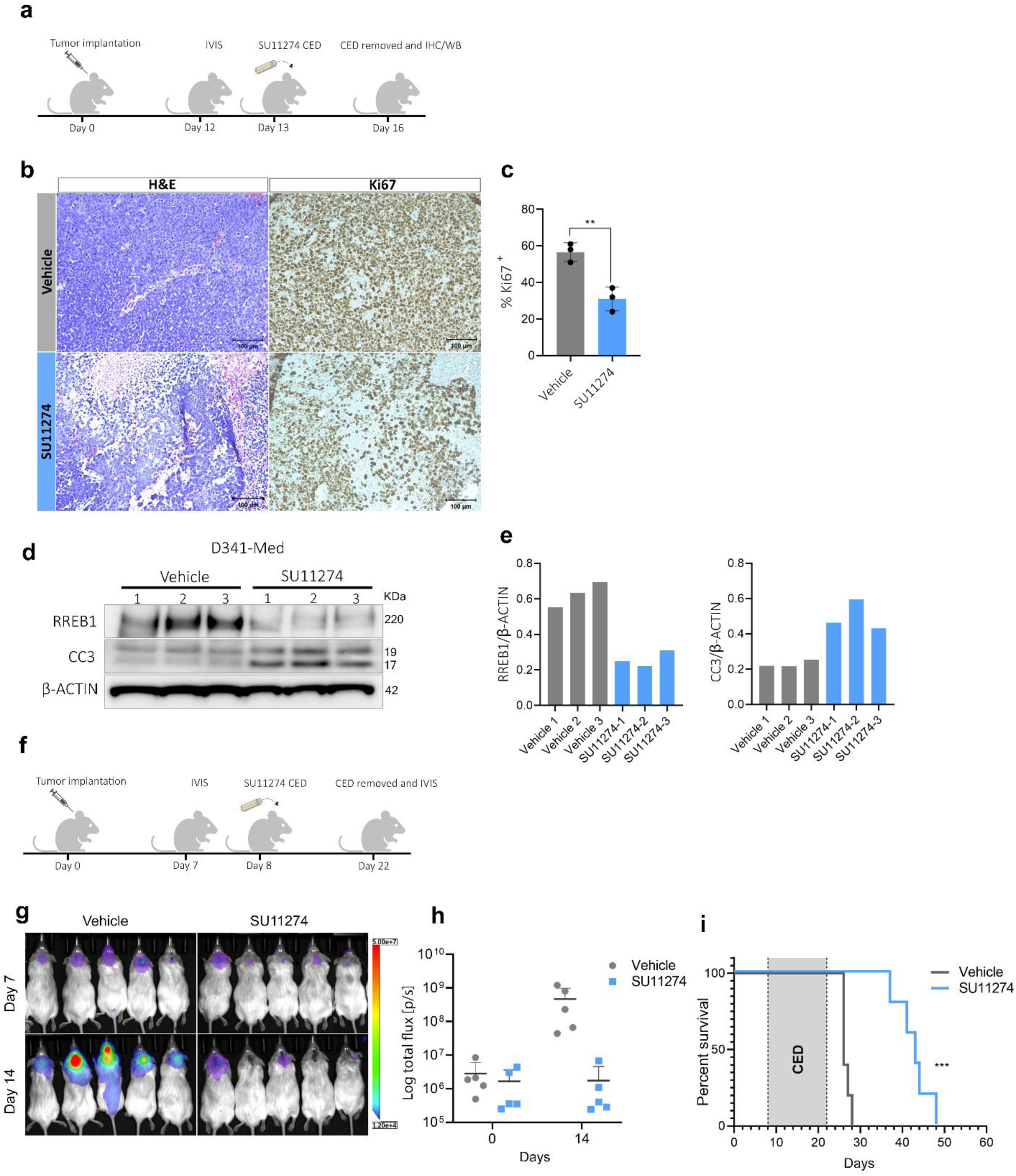
Inhibition of c-MET prolongs survival of G3-MB tumor bearing mice. **a.** Schematic timeline of short-term SU11274 administration in D341-Med tumor bearing mice. D341-Med cells were implanted and tumor growth was monitored on day 12 post-implantation. On day 13, osmotic pumps loaded with SU11274 (100 μM) or vehicle were implanted to deliver continuous drug infusion for 72-hours. Pumps were removed on day 16, at which point tumors were harvested for downstream analyses. **b.** Immunohistochemical detection of Hematoxylin and Eosin (H&E) and Ki67 in adjacent tumor sections. **c.** Quantification of Ki67 positivity was performed by scoring three independent fields of view. SU11274-treated tumors exhibited a significantly lower fraction of Ki67-positive nuclei (31 %) compared to vehicle controls (56 %). (Error bars represent mean ± SD, ** p < 0.01). **d, e.** Western blot analysis and densitometric quantification of RREB1 and cleaved caspase-3 (CC3) in D341-Med tumors following 3-day treatment with SU11274. Tumor-bearing mice received continuous delivery of SU11274 via osmotic pump for 72-hours, after which tumors were harvested and lysates subjected to SDS–PAGE and immunoblotting. Compared to vehicle-treated controls, SU11274-treated tumors exhibit decreased RREB1 protein levels alongside increased accumulation of CC3, indicative of enhanced apoptotic signaling. β-Actin serves as a loading control. **f.** Timeline of long-term SU11274 administration and survival monitoring in tumor-bearing mice. Tumor growth was assessed on day 7 post-implantation. On day 8, osmotic pumps delivering SU11274 (100 μM) or vehicle were implanted for 14 days. Pumps were removed on day 22, and tumors were imaged by IVIS. Thereafter, mice were monitored daily for survival and onset of neurological symptoms; animals reaching predefined humane endpoints were euthanized, and survival data were recorded. **g.** Bioluminescence imaging of NSG mice bearing medulloblastoma xenografts before (Day 0) and 14 days following continuous intratumoral delivery of 100 μM SU11274 or vehicle. **h.** Representative IVIS images demonstrate a pronounced reduction in tumor-associated luminescent signal in SU11274-treated mice relative to vehicle controls (n = 5 per group). Bioluminescence is expressed as Log10 total flux (photons/second) (n = 5 per group). **i.** Kaplan-Meier survival curves demonstrating prolonged median survival in SU11274-treated mice (43 days) compared to vehicle controls (26 days; log-rank Mantel-Cox test, *** p < 0.001).

To achieve prolonged drug administration for survival studies, we used osmotic pumps with a lower flow rate of 0.25 µL/hour, enabling a sustained two-week drug release. The pumps were loaded with SU11274 at a concentration of 100 µM and implanted orthotopically into the cerebellum of tumor-bearing mice (Fig. 6f). After two weeks, the pumps were removed, and tumor progression and survival were monitored over time until clinical endpoint symptoms were observed. Mice receiving intratumoral SU11274 exhibited a median survival of 43 days compared to 26 days in the vehicle treated group (Fig. 6g-i). These findings indicate that c-MET inhibition can effectively delay tumor progression in vivo and may represent a promising therapeutic strategy.

## Discussion

Medulloblastoma remains a significant clinical challenge, particularly in G3-MB patients, who exhibit the highest rates of metastasis and the worst prognosis. Our study identifies RREB1 as a transcription factor that is highly expressed in G3- and G4-MB, with minimal expression in WNT and SHH subgroups or normal brain tissue. Functional studies revealed that RREB1 is essential for tumor cell survival, as its knockdown significantly impairs proliferation and extends survival in tumor-bearing mice. Mechanistically, our data suggest that RREB1 regulates TGF-β target genes, further linking it to oncogenic pathways in MB. Additionally, we identified c-MET as a potential upstream regulator of RREB1, as pharmacologic inhibition of c-MET led to a reduction in RREB1 protein levels and prolonged survival in tumor bearing mice. These findings suggest that targeting c-MET could serve as a viable therapeutic approach for suppressing RREB1-driven oncogenic programs in these tumors.

Our studies demonstrate that RREB1 is essential for tumor proliferation and survival in G3-MB. Knockdown of RREB1 in G3-MB cell lines led to a significant reduction in cell viability and proliferation in vitro. Importantly, in vivo studies revealed that RREB1 knockdown extended survival in tumor-bearing mice, highlighting its functional significance in MB progression. However, overexpressing Rreb1 in combination with Myc in cerebellar neural stem cells was not sufficient to drive tumor growth, suggesting that additional oncogenic events are required for tumor initiation. Thus, RREB1 may function as a context-dependent oncogene, requiring cooperative interactions with other transcription factors or signaling pathways to fully exert its tumor-promoting effects.

Mechanistically, our RNA-seq and CUT&RUN analysis identified the TGF-β signaling pathway as a key target of RREB1. Knockdown of RREB1 resulted in downregulation of multiple TGF-β pathway genes, including *E2F5*, *ID1*, *INHBE*, and *GREM2*, with *E2F5* emerging as a crucial effector of RREB1-mediated oncogenic activity. Previous studies have shown that certain TGF-β receptors are amplified, while some TGF-β inhibitors are deleted in a subset of G3 tumors, supporting the notion that TGF-β signaling plays a complex role in MB (8). Recent work by Abeysundara et al. demonstrated that bone morphogenetic proteins (BMPs), key downstream effectors of the TGF-β superfamily, enhance metastatic colonization and tumor growth, further implicating the TGF-β signaling pathway in the promotion of aggressive disease phenotypes (22). Furthermore, other groups have demonstrated that pharmacologic inhibition of TGF-β signaling, such as with the small molecule galunisertib, can prolong survival in preclinical models of G3-MB (18). Our findings also align with recent reports that RREB1 can act as a cofactor for SMAD proteins, priming target gene enhancers for SMAD activation (23,24). RREB1 knockdown resulted in diminished SMAD-mediated transcriptional activity and downregulation of TGF-β target genes in both D283-Med and D341-Med cell lines. The consistent suppression of TGF-β target genes across both cell lines suggests that RREB1 primarily enhances SMAD target activation by modulating SMAD activity as suggested by reporter assay. Further investigation is warranted to elucidate the precise molecular interplay between RREB1 and SMADs that results in activating TGF-β-responsive gene networks.

Given RREB1’s role in MB pathogenesis, we sought to identify upstream regulators amenable to pharmacological targeting. A high-throughput kinase inhibitor screen revealed that the c-MET inhibitor SU11274 suppresses RREB1 protein levels. Notably, previous studies have implicated c-MET signaling in medulloblastoma progression, demonstrating that HGF can induce MYC expression in MB cell lines (25). In a separate study, inhibition of MET with PHA665752 effectively reduced the proliferative capacity of the D283, ONS76, and MED8A MB cell lines (26). Moreover, the selective MET inhibitor foretinib has been shown to suppress MET phosphorylation in SHH-subgroup MB cell lines, leading to pronounced decreases in proliferation and robust induction of apoptosis (27). Since SHH-MB expresses much lower levels of RREB1 than G3-MB, it seems likely that the effects of MET inhibition in SHH-MB are mediated through distinct mechanisms, independent of RREB1.

While the MAPK pathway has been previously linked to RREB1 regulation, our findings establish c-MET signaling as an additional critical modulator of RREB1 expression in MB. To evaluate the therapeutic potential of c-MET inhibition, we utilized CED to directly administer SU11274 into orthotopic MB tumors. This approach was necessary given the lack of evidence supporting SU11274’s ability to cross the blood-brain barrier. Short-term treatment with SU11274 resulted in decreased RREB1 and Ki67 levels and increased CC3 expression, indicating that c-MET inhibition promotes apoptosis and suppresses proliferation in vivo. Encouragingly, CED has already been successfully employed in clinical trials for central nervous system malignancies, demonstrating its potential as a therapeutic delivery platform (28–30). Additionally, while our study demonstrated that SU11274 effectively reduces tumor cell proliferation, it is unclear whether prolonged inhibition of c-MET may lead to resistance mechanisms that allow tumors to bypass RREB1 dependency. Future studies should examine potential pathways that may emerge following long-term c-MET inhibition.

In conclusion, our study provides strong evidence that RREB1 is a key driver of G3-MB and that its expression is maintained through c-MET signaling. Pharmacologic inhibition of c-MET using SU11274 effectively reduces RREB1 levels, impairs tumor cell proliferation, and extends survival in vivo. These findings establish a rationale for further preclinical and clinical investigation of c- MET inhibitors in MB treatment. Future studies to optimize drug delivery, refine combinatorial strategies, and validate these findings in clinically relevant models hold significant promise for advancing targeted therapies against G3-MB, ultimately offering hope for improved outcomes in this devastating pediatric malignancy.

## Methods

### Animal experiments

Mice were housed and bred under controlled environmental conditions at the Columbia University, Irving Cancer Research Center (ICRC) animal facility. All experimental procedures adhered to the NIH’s Guide for the Care and Use of Laboratory Animals and were approved by the Institutional Animal Care and Use Committee (IACUC) at Columbia University and Sanford Burnham Prebys Medical Discovery Institute. Mice used in this study were obtained from JAX Laboratory (NOD-SCID IL2Rgamma null, stock #005557).

Orthotopic tumor transplantation was performed by resuspending MB cell lines or patient-derived xenografts (PDXs) in VitroGel® 3D High Concentration (TheWell Bioscience, Monmouth Junction, NJ) and transplanting the mixture into the cerebellum of adult NSG mice via stereotaxic injection at the following coordinates: medial-lateral (ML: −1.0 mm), anteroposterior (AP: −7.0 mm), and dorsoventral (DV: −2.0 mm). Tumor progression was assessed via in vivo bioluminescence imaging (IVIS). Animals were euthanized upon reaching predefined humane endpoints based on neurological symptom severity, in accordance with IACUC guidelines.

### Cell culture

MB cell lines D283-Med, D341-Med, D425-Med, and DAOY were acquired from the American Type Culture Collection (ATCC) and Millipore Sigma (Cat# SCC290). HDMB03 cells were kindly provided by the laboratory of Dr. Tim Milde. For cell culture, D283-Med and DAOY cells were maintained in DMEM (supplemented with 4.5 g/L glucose, L-glutamine, sodium pyruvate), 10% FBS, 1× Pen/Strep, and 1× non-essential amino acids (NEAA). Similarly, D341-Med and D425-Med cells were cultured in identical DMEM base medium but with 20% FBS. HDMB03 cells were cultured in RPMI, 10% FBS, × L-glutamine, NEAA, sodium pyruvate, and 1× Pen/Strep. All cell lines were incubated under standard conditions (37°C, 5% CO₂).

### Lentiviral constructs

For overexpression of HA-tagged RREB1, the human RREB1 cDNA was subcloned from the Addgene plasmid pSPORT-RREB1 (#41145) into the LVX-IRES-GFP lentiviral vector (Takara Bio) using Gibson Assembly® (New England Biolabs). Lentiviral particles were generated in HEK293T cells via standard Polyethylenimine transfection, and target cells were transduced with 2 μg/mL polybrene (Santa Cruz Biotechnology, Cat# sc-134220). Constitutive and doxycycline-inducible shRNAs targeting RREB1 were obtained from Horizon Discovery (Dharmacon™) and E2F5 shRNA was obtained from Sigma-Aldrich (MISSION® shRNA, see Supplementary Table 2 for sequences). Lentiviral shRNA particles were produced and transduced as described above.

### Quantitative RT-PCR (qRT-PCR)

Total RNA was isolated from experimental cells using RNeasy Plus Mini Kit (Qiagen, Cat# 74134). and reverse-transcribed into cDNA using the iScript™ Reverse Transcription Supermix (Bio-Rad, Cat# 1708841) on a Bio-Rad DNA Engine® thermocycler. qPCR was performed in triplicate using the iQ™ SYBR® Green Supermix (Bio-Rad, Cat# 1708882) on a CFX384 Real-Time PCR Detection System (Bio-Rad). Primer pairs (sequences provided in Supplementary Table 2) were designed using Primer-BLAST (NCBI) and validated for efficiency. Relative gene expression was calculated using the ΔΔCt method, with β-ACTIN serving as the endogenous normalization control

### In vitro drug treatment

To assess protein-level changes in drug-treated cells in vitro, cells were serum-starved for 24-hours, followed by 48-hour treatment with the following compounds or ligands: SU11274 (MedChem Express, Cat# HY-12014), foretinib (MedChem Express, Cat# HY-10338), torkinib (MedChem Express, Cat# HY-10474), GNF-2 (MedChem Express, Cat# HY-11007), Asiatic acid (Selleck Chemicals, Cat# S2266), HGF (PeproTech, Cat# 100-39H).

### Cell viability and proliferation assays

Cell viability and proliferation were assessed using distinct experimental workflows. For viability assays, cells subjected to shRNA-mediated *RREB1* knockdown were plated in 96-well plates (20,000 cells/well), and viability was monitored over 10 days using the CellTiter-Glo® Luminescent Cell Viability Assay (Promega, Cat# G7570). Parallel experiments using E2F5 shRNA assessed viability over 8 days under identical conditions. For compound treatments, cells were exposed to HLM006474, SU11274 or foretinib for 72 hours prior to viability measurement. Reagent was added at a 1:1 (v/v) ratio to cultured cells, and luminescence was quantified using a Varioskan LUX Multimode Microplate Reader (ThermoFisher Scientific). For proliferation assays, cells treated with experimental compounds or shRNA were plated similarly and incubated for 72-hours. Proliferation was measured via BrdU incorporation using the BrdU Cell Proliferation Assay Kit (Cell Signaling Technology, Cat# 6813), following the manufacturer’s protocol.

### Flow cytometry

Infected cells expressing either RREB1 shRNAs or the shCtrl were sorted 48-hours post-infection and subsequently cultured for 5 days. Following this incubation period, the cells were stained with Annexin V (BD Pharmingen™, Cat# 556547) and 7-AAD (BD Pharmingen™, Cat# 559925) in Annexin V binding buffer (BD Pharmingen™, Cat# 556454) according to the manufacturer’s protocol. Stained cells were then analyzed using a BD LSRFortessa™ Cell Analyzer.

### CUT & RUN

D283-Med cells were transduced with a lentiviral construct expressing HA-tagged human RREB1. Transduced cells were expanded in culture, and aliquots of 1 × 10^6^ cells were processed for CUT&RUN or CUT&RUN-qPCR using the CUTANA™ CUT&RUN Kit, protocol v1.8 (EpiCypher, Durham, NC) or CUT&RUN Assay Kit (Cell Signaling Technology, #86652), respectively, per manufacturer protocols. The HA epitope tag was detected using the CUTANA™ HA-Tag antibody (EpiCypher, Cat# 13-2010) or Rabbit mAb IgG isotype control (Cell Signaling Technology, Cat# 3900S). Libraries were prepared with the NEBNext® Ultra™ II DNA Library Prep Kit for Illumina (E7645S), sequencing was performed by Novogene Co., Ltd. on the Illumina NovaSeq 6000 platform (PE150 configuration) with approximately 20 million paired-end reads per sample. For CUT&RUN-qPCR, the standard protocol was followed as above and qPCR was performed using gene specific primers (sequences provided in Supplementary Table 2)

### RNA sequencing

D283-Med cells were transfected with a doxycycline-inducible shRNA construct targeting RREB1 to achieve inducible knockdown. Protein knockdown was confirmed via immunoblotting at 24- and 48-hours post-induction, after which the 24-hour timepoint was selected for downstream analysis. Cells were treated with 2 μg/mL doxycycline to induce shRNA expression, harvested, and total RNA isolated using the RNeasy Plus Mini Kit (Qiagen, Cat# 74134).

Libraries were prepared with the TruSeq RNA Library Prep Kit v2. Sequencing was performed by Novogene Co., Ltd. on the Illumina NovaSeq 6000 platform, generating ∼20 million 150-bp paired-end reads per sample.

### Western Blotting

For Western blot analysis, cells were lysed in 1× RIPA buffer (Millipore, Cat# 20-188) supplemented with protease/phosphatase inhibitor cocktail (Cell Signaling Technology, Cat# 5872S). Total protein lysates were resolved on NuPAGE™ 3–8% Tris-Acetate gradient gels (Invitrogen, Cat# EA03752) and transferred to PVDF membranes (Invitrogen, Cat# LC2005). Membranes were blocked with 5% non-fat dry milk (Fisher Scientific, Cat# 50-488-785) in TBST (Tris-buffered saline with 0.1% Tween-20) for 1 hour and probed overnight at 4°C with the following primary antibodies: RREB1 (1:1000, Sigma-Aldrich, Cat# HPA001756); HiBiT (1:1000, Promega, Cat# N7200), SMAD2/3 (1:1000, Cat# 8685S), phospho-SMAD2 (1:1000, Cat# 3108T), ERK1/2 (1:1000, Cat# 9102S), phospho-ERK1/2 (1:1000, Cat# 9101S), AKT (1:1000, Cat# 9272S), phospho-AKT (1:1000, Cat# 4051S), Cleaved-Caspase 3 (1:1000, Cat# 9664), β-actin (1:1000, Cat# 4970S), and GAPDH (1:1000, Cat# 2118S) (all Cell Signaling Technology unless noted). Following primary antibody incubation, membranes were treated for 1 hour with HRP-conjugated anti-rabbit (Cat# 7074S) or anti-mouse (Cat# 7076S) secondary antibodies (1:1000, Cell Signaling Technology). Protein signals were detected using Clarity™ Western ECL Substrate (Bio-Rad, Cat# 170-5060) and imaged on a ChemiDoc™ MP Imaging System (Bio-Rad).

### SMAD Reporter assay

To assess SMAD2/3/4 transcriptional activity following RREB1 knockdown, stable D341-Med cells were generated by lentiviral transduction with a SMAD2/3/4-responsive luciferase reporter construct (System Biosciences, Cat# TR203PA-P). Cells were transduced with RREB1-targeting shRNA for 72 hours, with or without recombinant human Activin B (200 ng/ml, R&D Systems, Cat# 11517-AB-010) or TGFβ (200 ng/ml, PeproTech, Cat# 100-21) after which reporter activity was quantified. Briefly, 150 μg/mL D-luciferin (PerkinElmer, Cat# 122799) was added to cells, followed by a 2-minute incubation at room temperature. Luminescence signals were measured using a Varioskan LUX Multimode Microplate Reader (Thermo Fisher Scientific).

### CRISPR-Cas9 knock-In

A HiBiT-tag sequence (Promega) was synthesized by Integrated DNA Technologies (IDT) and integrated into the N-terminus of the human RREB1 locus via CRISPR-Cas9 genome editing. A single-guide RNA (sgRNA) targeting RREB1 was designed, and ribonucleoprotein (RNP) complexes were assembled by incubating 8 μg of TrueCut™ Cas9 Protein v2 (Invitrogen), 50 μM sgRNA, and 70 μg of HiBiT donor template in Buffer GE (Invitrogen) for 15 minutes at room temperature. The RNP complexes were electroporated into 2 × 10^5^ HDMB03 cells using the Neon™ NxT Electroporation System (Invitrogen) with parameters set to 1000 V, 40 ms, and 1 pulse. Electroporated cells were recovered in pre-warmed culture medium, expanded, and single-cell sorted into 96-well plates. Clonal populations were screened by Sanger sequencing to confirm HiBiT-tag knock-in. HiBiT-tagged cells were plated in 96-well plates, and luminescent signals were quantified using the Nano-Glo® HiBiT Lytic Detection System (Promega, Cat# N3030) following the manufacturer’s protocol.

### High-throughput drug screening

Cells were seeded into 384-well tissue culture-treated microplates (Greiner Bio-One, Cat# 781080) at a density of 5,000 cells per well in appropriate growth medium. The following day, compounds from the SelleckChem kinase inhibitor library, comprising 378 small-molecule inhibitors, were dispensed using an Echo 550 acoustic liquid handler (Beckman Coulter). A 25 nl volume was transferred per well to achieve a final drug concentration of 10 µM. Nano-Glo® HiBiT lytic detection (Promega) was added to the wells at 4 and 8-hours post-treatment. Luminescence was measured using an EnVision Multimode Plate Reader (Revvity) according to the manufacturer’s protocol.

### CED surgeries

Tumor-bearing mice were anesthetized, and 2-week CED osmotic pumps (ALZET, Cat# 1002) were implanted subcutaneously. A cannula was stereotactically positioned at the site of prior tumor implantation using a Brain Infusion kit 3 (ALZET, Cat# 0008851), adjusted to a depth of 2 mm. The cannula was secured to the skull with Loctite adhesive (ALZET, Cat# 0008670) and reinforced with dental cement (Fisher Scientific, Cat# 10-000-786). Pumps delivered 100 µM SU11274 (or vehicle control) prepared in artificial cerebrospinal fluid (aCSF; recipe per ALZET guidelines) containing 1% DMSO and 20% PEG300, at a flow rate of 0.25 µL/hour. After 14 days, pumps were surgically removed, and tumor progression was assessed via bioluminescence imaging using an IVIS® Spectrum system (PerkinElmer).

### Immunohistochemistry

Paraffin-embedded tissue sections were deparaffinized and rehydrated through graded alcohols. Antigen retrieval was performed using a heated citrate-based solution (Vector Laboratories, Cat# H-3300-250) under pressurized conditions. Endogenous peroxidase activity was quenched by incubating the sections in 3% hydrogen peroxide (Millipore Sigma, Cat# 1.07209) for 10 minutes. Non-specific binding was blocked with 2.5% horse serum (Vector Laboratories, Cat# S-2000-20) for 1 hour at room temperature. Sections were then incubated overnight at 4°C with an anti-Ki67 primary antibody (1:200 dilution; Cell Signaling Technology, Cat# 9449T). Following washes in TBST, sections were incubated with an ImmPRESS™ horse anti-mouse IgG secondary antibody (Vector Laboratories, Cat# MP-7402-NB) for 30 minutes at room temperature. Signal detection was carried out using DAB chromogen (Vector Laboratories, Cat# SK-4105), and counterstaining was performed with a 1:10 dilution of hematoxylin (Vector Laboratories, Cat# H-3404-100).

Finally, sections were mounted using a permanent mounting medium (Vector Laboratories, Cat# H-5501-60), and images were acquired using a Zeiss LSM 700 microscope (Carl Zeiss, Jena, Germany). Ki67 staining was quantified using imageJ (31) with Color Deconvolution2 plugin.

### Bioinformatic analysis

Comparative analysis of transcription factors:

Processed and normalized gene expression datasets from Cavalli et al. (2017; GSE85217) were retrieved from the R2 Genomics Analysis and Visualization Platform (https://hgserver1.amc.nl/). Differential gene expression was conducted using a |log2 fold change| (FC) > 1 and false discovery rate (fdr) threshold of < 0.01. Genes were filtered based on their annotated DNA-binding transcription factor activity according to the Kyoto Encyclopedia of Genes and Genomes (KEGG) database. Volcano plots were generated using the VolcaNoseR2 platform (https://huygens.science.uva.nl/VolcaNoseR2) (32), applying a |log2FC| > 0.6 and an fdr < 0.01 for visualization.

### Proteomics data

Proteomic expression profiling was performed on the Ayrault lab cohort, comprising 343 primary medulloblastoma samples, using mass spectrometry as previously described (manuscript in preparation). Subgroup and subtype classification was based on the DNA methylation-based framework for central nervous system tumors (33). Briefly, the mass spectrometer was operated in data-independent analysis (DIA) mode for peptide acquisition. Raw data files were analyzed using Spectronaut 19 with the spectral library build in-house and further processed and normalized using myProMS v3.10 (https://github.com/bioinfo-pf-curie/myproms) (34). The accession number for the proteome raw dataset will be available in PRIDE repository once the Ayrault’s paper is published.

### Methylation analysis

MB tumors and postnatal CB samples with matched DNA methylation and gene expression profiles were obtained from previously published datasets (14,15). DNA methylation data were generated using Illumina 450k arrays and processed with the minfi R package (v1.18.0) employing the default Illumina preprocessing function to derive CpG beta values. Gene expression data were derived from raw intensities of Affymetrix U133 Plus 2.0 arrays and normalized using the affy R package with MAS5 normalization.

### RNA-seq

Subread 2.0.3 (35) was used to align the reads to the GRCh38 build of the human genome. featureCounts (36) was used to quantify the expression of each gene. Three methods were used to check for quality and outliers: Multidimensional scaling (37), as implemented in limma 3.5.62 (38); Hierarchical clustering (39) as implemented in gplots 3.1.3 (40); and Principal Component Analysis (41) as implemented in affycoretools 1.72.0 (42). Differential gene expression was analyzed with limma-voom with sample weighting (38,43,44).

The SIPA (Signal Interaction Pathway Analysis) (45,46) method, as implemented in iPathwayGuide iPathwayGuide (47), was used to perform KEGG (48) analysis. Differential gene expression cutoffs for pathway analysis were of p-value ≤ 0.001 and |log2 fold change| ≥0.6. High specificity pruning, an enhanced version of the elim method (49), implemented in iPathwayGuide, was used to perform Biological Process Gene Ontology (GO) (50) analysis. Webgestalt was used to analyze the data in terms of Reactome (51) using overrepresentation analysis (52).

### CUT&RUN

Bowtie2 (53) was used to align the reads to the GRCh38 build of the human genome, using parameters were that are recommend for this analysis by the developers of CUT&RUN (54). Removal of duplicates and sorting of sam files was performed using a script furnished by the developers of SEACR (55). Differential binding sites were identified with the SEACR online tool (56). Intersection of binding sites from replicate samples was performed using BEDTools (57).

BEDTools was also used for file format transformation, except where noted. The location of nearest genes to each peak was determined with ChIPpeakAnno (58). Both the nearest TSS to a binding site, and all genes that overlapped with the binding site were detected. Pathways overrepresented by detected sites, using rGREAT (59), which is the R implementation of the GREAT (Genomic Regions Enrichment of Annotations Tool) method (60). Coverage plots were generated with scripts with commands from bedtools, deepTools2 (61,62), and bedGraphtoBigwig. Integrated genomics viewer was used to display gene-specific coverage plots. deeptools2 was used to display scaled genome-wide coverage plots.

Integration of CUT&RUN with RNA-seq:

Differentially expressed genes associated with pathways of interest were found using Venn operations in R. Sequences near genes of interest were obtained using BSgenome 1.68.0 (63).

## Data availability

The RNA-sequencing data generated and analyzed during this study have been deposited in the NCBI Gene Expression Omnibus (GEO) under the accession number GSE298167. The CUT&RUN sequencing data have been deposited in GEO under the accession number GSE298168. All datasets are publicly available and can be accessed at https://www.ncbi.nlm.nih.gov/geo/.

## Acknowledgements

This work was supported by R35 NS122339 from the National Institute for Neurological Disorders and Stroke (RJWR), R01 CA159859 (RJWR and MDT), William’s Superhero Fund, and by McDowell Charity Trust. The authors gratefully acknowledge the Animal Care Program at the University of California, San Diego, and the Genomics Shared Resources at Sanford Burnham Prebys Medical Discovery Institute (supported by P30 CA030199), as well as the Biomedical Informatics, OPTIC, and Histology Core Shared Resources in the Herbert Irving Comprehensive Cancer Center at Columbia University (supported by P30 CA013696). We would like to thank Anindya Bagchi, David Largaespada, Theophilos Tzaridis, and Tanja Eisemann for thoughtful discussions.

## Author Contributions

Conceptualization: RJWR, MBM.

Methodology: RJWR, MBM, RAF, SP, BLG, JTD, LQC, SG, YL, OC, PS, LG.

Investigation: MBM, KRC, CL, GF Supervision: RJWR, PAN, AR, LC, MDT Corresponding author

Correspondence to Robert J. Wechsler-Reya.

## Ethics declarations

### Competing interests

The authors declare no competing interests.

## Notes

### Competing Interest Statement

The authors have declared no competing interest.

