## Supplementary figures for "Ras-Responsive Element Binding Protein 1 regulates survival of Group 3 medulloblastoma"

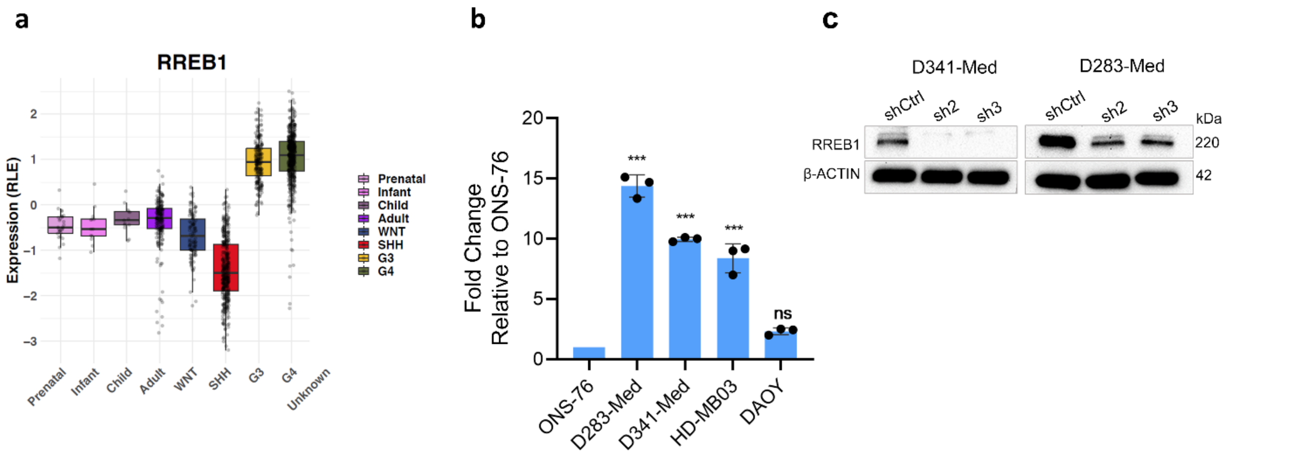


**Supplementary Figure 1.**

**a.** RREB1 mRNA expression levels in healthy prenatal, infant, child, adult and subgroups of MB. Data is acquired from *Weishaupt et al, 2019*, comprised of 1350 MB and 291 normal brain samples (RLE: Relative log expression). **b.** qPCR RREB1 mRNA expression in MB cell lines measured by ΔΔCt method. Error bars represent mean ± SD from three biological replicates (*** $p < 0.001$, ns = non-significant). **c.** Western blotting showing RREB1 knockdown efficiency 48-hours post-transduction.

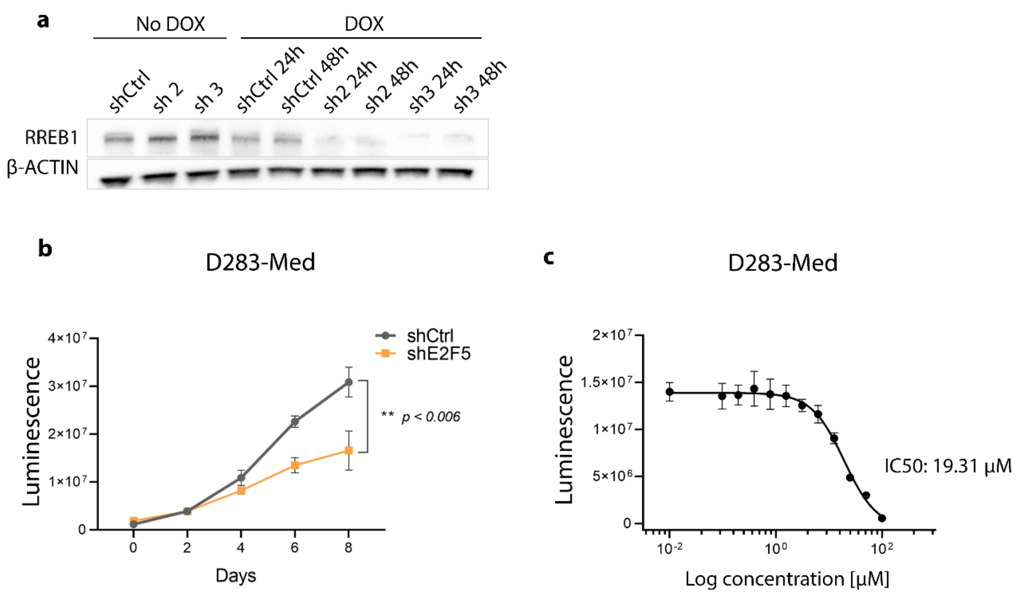


**Supplementary Figure 2.**

**a.** Validation of RREB1 protein knockdown using a doxycycline-inducible shRNA system. Cells were transduced with either a non-targeting control shRNA (shCtrl) or an shRNA targeting RREB1, followed by selection with 2 µg/mL puromycin. Doxycycline was added to induce shRNA expression for 24 and 48-hours, after which RREB1 protein levels were assessed by Western blotting. **b.** Cell viability of D283-Med cells following E2F5 knockdown, assessed using the CellTiter-Glo assay. Statistical significance was determined by unpaired *t*-test; ** $p < 0.01$. **c.** Dose-response curve of D283-Med cells treated with the E2F5 inhibitor HLM006474. Cells were treated for 72-hours, resulting in an IC50 of 19.31 µM. Data represent mean ± SD from three independent experiments (n = 3).

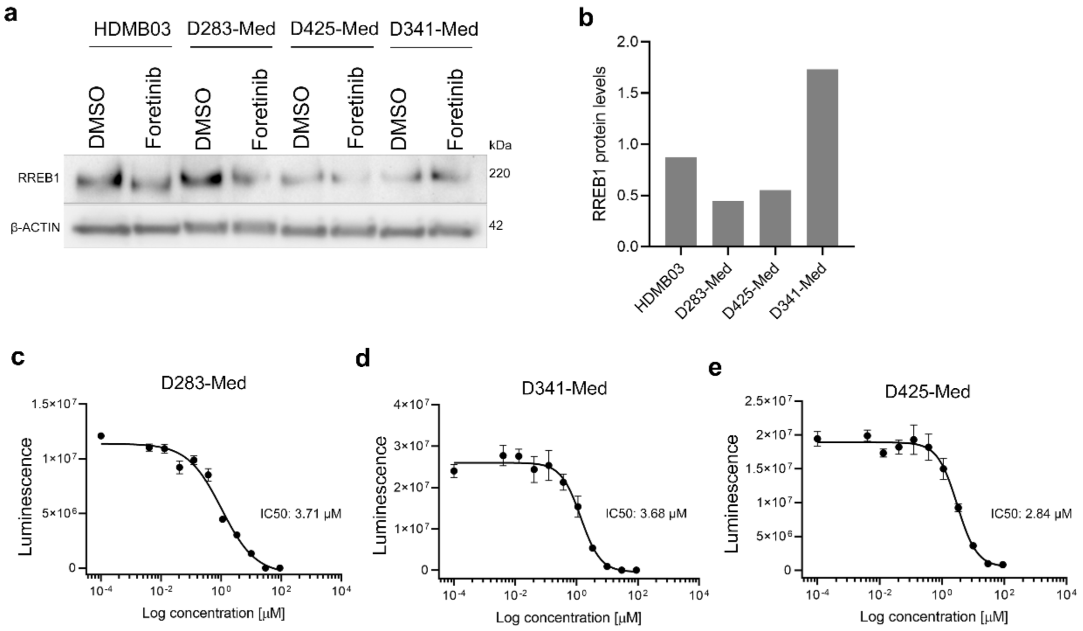


**Supplementary Figure 3.**

**a, b.** Foretinib treatment reduces RREB1 protein levels in medulloblastoma (MB) cell lines. Western blot analysis of RREB1 protein following treatment with 3 µM foretinib for 48 hours. **c-e.** Dose-response curves of MB cell lines treated with foretinib for 72 hours, assessed using the CellTiter-Glo assay. Calculated IC50 values: D283-Med, 3.71 µM; D341-Med, 3.68 µM; D425-Med, 2.84 µM. Data represent mean ± SD from three independent experiments (n = 3).

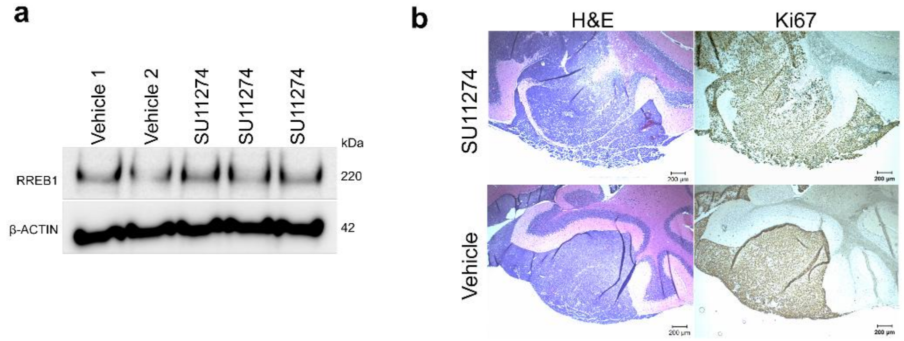


**Supplementary Figure 4.**

**a.** RREB1 protein levels in medulloblastoma tumors following systemic administration of SU11274. Tumor-bearing mice were treated by oral gavage with SU11274 (10 mg/kg) for 3 days, and RREB1 expression was assessed by Western blotting. **b.** Low magnification (5x) images of Hematoxylin and Eosin (H&E) staining and Ki67 immunohistochemistry of MB tumors following 3 days of treatment with SU11274 delivered via osmotic pumps. Pumps were removed after 3 days, and brains were harvested and fixed for histological analysis.
